# Hormone levels increase in response to video playback in the Siamese fighting fish

**DOI:** 10.1101/2022.02.20.481239

**Authors:** Deepa Alex, Sara D. Cardoso, Andreia Ramos, David Gonçalves

**Author notes:** Corresponding author; USJ, Rua de Londres 106, Macao SAR. Correspondence: Deepa Alex,; Andreia Ramos,; Sara Cardoso,; David Gonçalves,.

## Abstract

The mechanisms of action of hormones regulating aggression in fish remain poorly understood. One possibly confounding variable is the lack of standardization in the type of stimuli used to elicit aggression. The presentation of controlled stimuli in videos, a.k.a. video playback, can provide better control of the fight components. However, this technique has produced conflicting results in animal behaviour studies and needs to be carefully validated. Here, the response of male *Betta splendens* to a matched for size conspecific fighting behind a one-way mirror presented live or through video playback was compared. Despite their non-interactive nature, both stimuli elicited a strong aggressive response by focal fish, which started with stereotypical threat displays and progressed to overt attacks. Overall, the frequency and duration of aggressive behaviours and swimming activity were similar towards live and video stimuli. Post-fight plasma levels of the androgen 11-ketotestosterone were elevated as compared to controls, regardless of the type of stimuli. Cortisol also increased in response to the video images, as previously described for interactive live fights in this species. The study shows for the first time in a fish a robust endocrine response to video stimuli and supports the use of this technique for researching aggressive behaviour in *B. splendens*.

## Introduction

Hormones are thought to explain a significant part of the variation in aggression levels both within and across species [1]. Steroid hormones, in particular androgens and corticosteroids, have been identified as relevant modulators of aggression but significant variation across studies exists. A conceptual framework to explain the role of androgens on the regulation of aggression was proposed in 1990 by John Wingfield and collaborators, on what became known as the challenge hypothesis [4]. Under this hypothesis, androgen levels are modulated by the conditions of the social environment, with animals living in more unstable contexts increasing circulating androgens above reproductive levels to facilitate the expression of mating and territorial behaviour. In fish, a meta-analysis has generally confirmed the assumptions of the challenge hypothesis [5], but contradictory evidence also exists in the literature [6–8]. A similar scenario occurs for the role of corticosteroids as modulators of fish aggression. In general, it has been shown that individuals with low basal levels of cortisol (F) are more aggressive [9] and that exogenous F administration reduces aggression [10]. However, other studies have shown that plasma F levels increase after fights [11,12], questioning the role of this hormone in the modulation of fish aggression.

These contradictory results may partially be a consequence of a lack of standardization in experimental procedures between studies. Different types of aggression-eliciting stimuli have been used when investigating aggressive behaviour in fish (for a review in zebrafish see [13]). These include paired fights with fish placed in the same arena (e.g., [14]), an opponent presented behind a transparent partition (e.g., [15]) or a one-way mirror (e.g., [16]), mirror (e.g., [16]) and nonreversed mirror images (e.g., [17]), static 2D or 3D models (e.g., [18]), robots (e.g., [19]), or video playback, i.e. the presentation of stimuli in video (e.g., [20]). Some of these stimuli are interactive and feedback received varies according to the behaviour displayed by the focal fish (e.g., opponents in the same arena). Mirror images provide interactive and symmetric stimuli as the feedback received is the same as the behaviour displayed by the focal animal. Others are non-interactive (e.g., one-way-mirrors) and may further be invariant across trials (e.g., video playback). The choice of the type of test to be used depends on the characteristics of the species under study and the objectives of the work. While live fights, where opponents are placed in the same arena, allow a resolution of the challenge, with a winner and a loser being determined, this is more difficult or impossible to achieve with other types of tests. However, it may not be ethical in highly aggressive species to conduct live fights as animals can get injured or even die. Further, if what is intended is comparing aggression levels across experimental groups, it may be more adequate to present animals with standardized stimuli that do not vary with the behaviour of the focal animal. Static models have been used for this purpose but they lack motion and may not be perceived as meaningful stimuli [13]. A few studies in fish have used robots and this is a promising technique as it allows a 3D interaction with a performing stimulus [19]. However, to build a robot that is recognized as a conspecific is complex and many labs lack the human resources and equipment to implement this technique. The presentation of standardized stimuli in a video to fish has been used for several decades with variable levels of success. While some species respond to video playback, others do not seem to recognize images in videos or present only a weak response, and this variation is thought to be related to sensorial differences within and across species (for a discussion see [21] and other articles in that volume). A commonality to the use of artificial stimuli to study behaviour is thus the need to properly validate that they are being perceived meaningfully. For this, equivalent artificial and natural stimuli should be used for comparison, which is not always the case (e.g., [22]). Further, while behaviour is usually measured, no study so far has used physiological endpoints to compare the response to video playback and live stimuli in fish. This is relevant because it has been shown that the physiological response to different stimuli may differ, even when the behavioural output does not [23].

Here, we compared the behavioural and endocrine response of male Siamese fighting fish *Betta splendens* to video playback of a conspecific and one-way mirror fights. For the one-way mirror test, a conspecific was observed by the focal fish fighting behind the one-way mirror. This provided a non-interactive fight as the conspecific stimulus could not observe the focal fish but rather its own image in the reflective side of the mirror. For the video playback trials, conspecific males fighting the one-way mirror were filmed from the same perspective of the focal fish and played back, providing an equivalent stimulus to the live opponent. Aggressive behaviour and post-fight androgen (11-ketotestosterone, KT) and corticosteroid (F) plasma levels were measured.

Overall, the study aimed to test and validate the use of video playback to investigate the endocrine regulation of aggressive behaviour in *B. splendens* and to provide additional data on post-fight endocrine responses to aggression in this species.

## Methods

### Animals

Siamese fighting fish used in this experiment were 17-month-old from the F1 generation of a cross between a wild-type and a fighter strain raised in mixed-sex groups (see [11] for more details on this line). Thirty-six males were isolated at ten months old into 9W X 9D X 20H cm tanks, containing a small ceramic shelter, with no visual contact with other conspecifics. Fish were fed twice a day, except on the day of the experiment where they were not fed, and maintained under a 12L:12D photoperiod and water temperature of 28 ± 1 °C. Reverse osmosis (RO) water for stock and isolation tanks was conditioned with Indian almond tree leaves and salinity was kept at 1 g/L. Two months prior to the beginning of the experiment, fish were transferred to new individual tanks of the same dimensions as the experimental tanks (25 W X 12.5 D X 20 H cm). Fish were netted and released back into their tank daily for one week before the experiment to habituate to handling.

### Experimental procedure

Fish were randomly assigned to one of four treatments: one-way mirror conspecific (*n* = 8), one-way mirror conspecific control (*n* = 7), video playback (*n* = 10), and video playback control (*n* = 7). Between groups, fish did not differ in weight (W) or standard length (SL) (One-way ANOVA, W, F_(3,32)_ = 1.006, p = 0.403; SL, F_(3,32)_ = 1.118, p = 0.356). Fish used as stimuli, both live and in video playback, and as focal were matched for size (SL, C.V. < 10%). The experiment was performed in an arena with 122 W X 57 D X 57 H cm, composed of white walls and two white sliding doors. For the one-way mirror setup, the tank of the focal fish was separated from the tank of the conspecific by a 1 cm thick one-way mirror (Fig. 1A). Two opaque smart screens that become transparent when activated prevented the focal fish from seeing the stimulus tank side and the stimulus fish from seeing the mirror during the acclimation period. The tank of the stimulus conspecific was narrower than the focal fish tank to avoid large variation in target size, with dimensions of 12.5 W X 8 D X 20 H cm. This tank had directly above it a diffuse LED strip to create the light contrast needed for the one-way mirror to be reflective for the stimulus but not for the focal fish. A diffuse LED panel provided general illumination to the arena. The behaviour of the focal fish was recorded using one side and one top Raspberry Pi camera module V2, and of the stimuli conspecific fish using a similar side camera, all at a resolution of 1640 × 922 px at 30 fps. Each camera was connected to an independent Raspberry Pi board 4B. Stimuli fish were used only once. For the one-way mirror control, no fish was added as stimulus and the focal fish observed an empty tank. Control trials were run with only the focal or stimulus fish added to the setup, ensuring that the focal fish was not viewing its mirror image and that the stimulus fish was responding to its mirror image and not to the focal fish. For the video playback trials, conditions were similar but an LCD screen (2560×1600 IPS) controlled remotely using a Raspberry Pi board 4B was placed adjacent to the focal fish tank (Fig. 1B).

**Figure 1.**
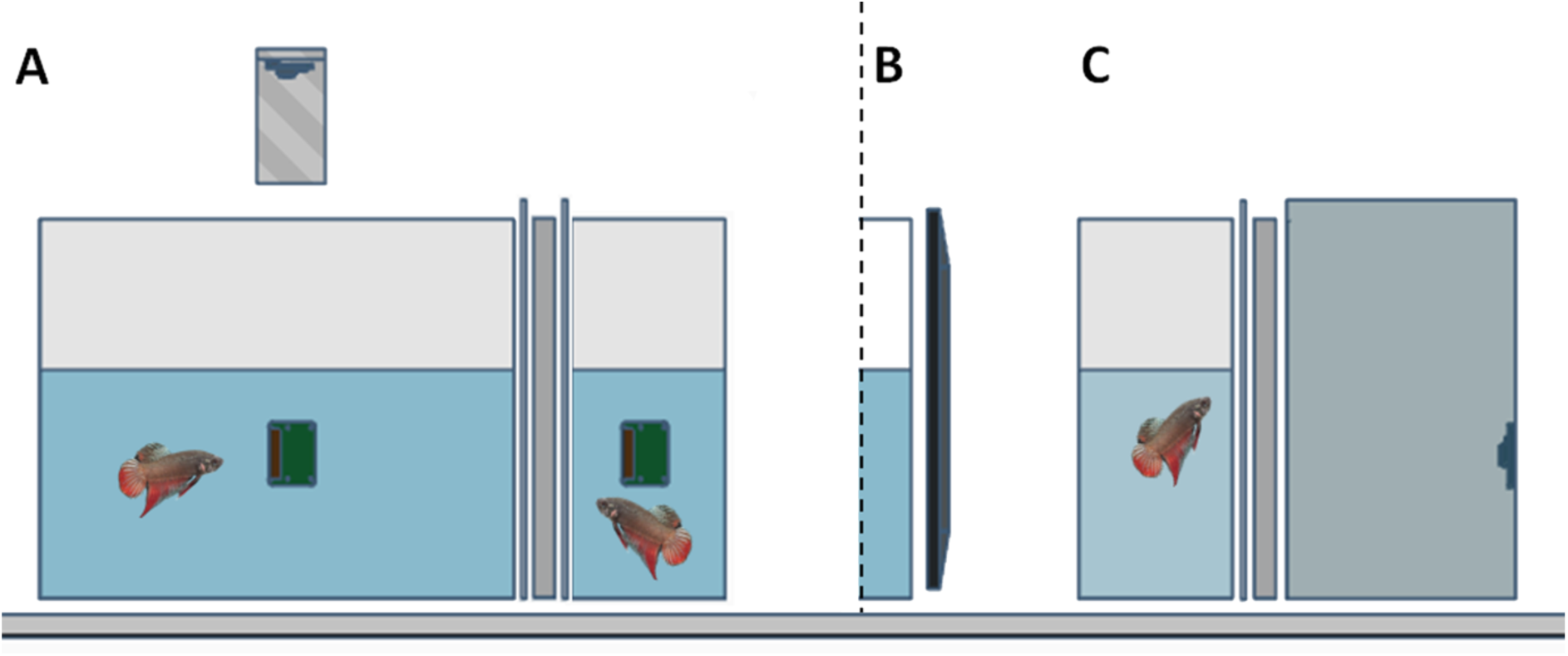
Experimental setup used to test (A) the response to a live conspecific fighting its mirror image or (B) to a video playback of an equivalent stimulus; (C) footage for the video playback was obtained by filming behind a one-way mirror a fish fighting its image. An opaque smart screen that becomes transparent when activated was placed between the one-way mirror and both the focal and the live stimulus tank and turned on after the acclimation period. Trials were recorded with top and side cameras for behavioural analysis.

Recording of the stimuli to be played back was done with a Logitech HD Pro Webcam C920 placed inside a dark chamber facing the one-way mirror and stimulus tank, separated by a smart screen, at a resolution of 1920 × 1080 px and framerate of 30 fps (Supplementary video 1). The stimulus fish was placed inside the tank and left to acclimate for 30 min. At the end of this period, the smart screen and video camera were activated, with recordings lasting 30 min. This allowed obtaining 30 min footage of a fish fighting its mirror image from the opponent’s perspective. The size of the fish on screen was 1:1 when the fish was displaying close to the mirror. A total of 7 fish were recorded and because of the need to match the stimulus and focal fish for size, one video was presented to three focal, another to two focal and the remaining were only used once. For the video playback control treatment, footage of the tank without any fish was obtained under the same conditions. Brightness of the LED screen was visually adjusted to match the tank light conditions as seen through the one-way mirror.

To begin a trial, fish were transferred from their housing tanks to the test tank, the cameras were activated, and a 30 min acclimation period was given with the smart screens opaque (one-way mirror trials) or a white screen on the LCD (video playback trials). At minute 30, the smart screens or the test video in the LCD screen were automatically started allowing the fish to see either: a live conspecific fighting its mirror image, an empty tank, a video of a conspecific fighting its mirror image, or a video of an empty tank. Observations had a total duration of 30 min. At the end of the observation, the focal fish was immediately removed from the tank, anaesthetized with cold buffered MS222 (concentration 600 mg/L) and blood extracted from the caudal vein using a heparinized 27G syringe. Time for blood extraction since the end of the trial was not recorded but in other similar experiments in our lab blood is collected within 2-5 min. After the procedure, individuals were placed in individual recovering tanks with aeration. Blood samples were centrifuged for 15 minutes, and plasma was transferred to new tubes and stored at −20°C until further analysis. Water in the test tank was changed between experimental trials.

### Behavioural analysis

Aggressive behaviours of both the focal and stimulus fish were manually scored with Boris software (V. 7.9.19 for Mac, [24]) and included: duration of open opercular displays, distended fins, and darkened body colour, and frequency of caudal swings, charges, bites, and air breathing. Total distance moved, time spent near the stimuli (within 5 cm) and near the surface of the tank (within 4 cm) was measured using Ethovision XT.

### Hormone analysis

Plasma levels of KT and F were measured with competitive enzyme-linked immunosorbent assay (ELISA) kits from Cayman Chemical following the manufacturer’s instructions. A lack of interference in the assay of other immunoreactive molecules for this species and for these ELISA kits had already been confirmed by serially diluting a plasma pool and comparing the slope with that of a standard curve [11]. All standards and samples were measured in duplicate with a dilution in the EIA buffer of 1:150 for KT and 1:20 for F. Experimental samples were measured in the same assay and the intra-assay coefficient of variation, calculated from the sample duplicates, was 2.86 % for KT and 3.20 % for F.

### Statistical analysis

Parametric procedures were used for all comparisons, with normality and homoscedasticity of data being tested a priori with Shapiro-Wilk’s and Levene’s tests, respectively. As indicated, some variables were log-transformed to comply with parametric assumptions or non-parametric tests were used when assumptions were still violated after data transformation. First, differences between the two control groups in total distance travelled, frequency of air breathing, time spent close to the surface, time spent close to the stimulus tank/screen side and hormone levels (KT and F) were analysed with unpaired t-tests. As expected, there were no differences for any of the variables between these two groups (P > 0.212) and they were merged for further analysis. For these same variables, differences between the experimental and the merged control group were tested with one-way ANOVAs with factor treatment (one-way mirror, video playback, control), followed by post-hoc Tukey tests to assess differences between groups. Differences in aggressive behaviours between the one-way mirror and video playback groups were tested with unpaired t-tests or with Mann-Whitney U tests as aggression was not displayed during control trials. All correlations between variables were assessed with Pearson’s correlation tests.

A principal component analysis (PCA) that included all measured behavioural and endocrine variables was applied to investigate the distribution of fish from each treatment. Variables were standardized prior to the analysis. It was predicted that control fish should form one cluster and that fish exposed to the conspecific, live or through video playback, should form another cluster if video playback is equivalent to a live stimulus, or two further clusters otherwise. Average distance to centroid in the two first PCA factors were calculated with package “vegan 2.5-7” for R.

All statistical analyses were run with R 4.1.0 [25].

### Ethics statement

Experiments followed the ASAB/ABS “Guidelines for the treatment of animals in behavioural research and teaching” [26]. The study complied with the ethical guidelines enforced at the University of Saint Joseph and experimental research with this species was approved by the Division of Animal Control and Inspection of the Civic and Municipal Affairs Bureau of Macao, license AL017/DICV/SIS/2016.

## Results

### Behaviour

Focal fish exhibited a clear aggressive response to both the live and video playback stimulus (Supplementary video 1). Activity and position related variables and metabolic effort, as determined from the frequency of air breathing [27], did not differ between these two treatments (Table 1). Likewise, the frequency or duration of aggressive displays and the time spent with aggressive colouration was similar (Table 2).

**Table 1.** Descriptive statistics, main effects and post-hoc comparisons of variables related with activity and position in the tank of fish presented with an empty tank or white screen (control), opponent behind a one-way mirror in an adjacent tank, and video playback of an opponent fighting a one-way mirror.

| | Control<br>Mean ± S.E. | One-way mirror<br>Mean ± S.E. | Video playback<br>Mean ± S.E. | One-way ANOVA main effects | | | Post-hoc comparisons - p | | |
| --- | --- | --- | --- | --- | --- | --- | --- | --- | --- |
|  |  |  |  | F | d.f. | p | Control vs<br>one-way<br>mirror | Control vs<br>video<br>playback | One-way mirror<br>vs<br>video playback |
| **Total distance travelled (cm)\*** | 1533.61 ± 123.01 | 2100.16 ± 130.06 | 2588.70 ± 379.39 | 5.444 | 2, 29 | **0.010** | 0.092 | **0.011** | 0.765 |
| **Time close to stimuli (s)\*** | 687.00 ± 42.41 | 1474.59 ± 133.57 | 1181.83 ± 129.84 | 18.464 | 2, 29 | **< 0.001** | **< 0.001** | **< 0.001** | 0.215 |
| **Time close to surface(s)** | 752.87 ± 64.70 | 753.52 ± 105.20 | 715.83 ± 81.29 | 0.07 | 2, 29 | 0.932 | 1.000 | 0.937 | 0.950 |
| **Frequency of air breathing** | 23.00 ± 2.62 | 31.62 ± 7.87 | 40.50 ± 7.02 | 2.845 | 2, 29 | 0.074 | 0.525 | 0.061 | 0.550 |

\*, log-transformed variables; statistically significant values are in bold.

**Table 2.** Descriptive statistics and paired tests of aggressive behaviours of focal fish in response to an opponent fighting a one-way mirror in an adjacent tank, presented live or by video playback. Fish from the control group did not exhibit any of these behaviours.

|  | One-way mirror |  | Video playback |  | T-test / Mann-Whitney U |  |  |
| --- | --- | --- | --- | --- | --- | --- | --- |
| | N | Mean ± S.E. | N | Mean ± S.E. | t / U | d.f. | p |
| Time with open opercula (s) | 8 | 210.83 ± 37.16 | 10 | 222.33 ± 47.69 | 0.183 | 16 | 0.857 |
| Time with fins distended (s)* | 8 | 1775.35 ± 12.64 | 10 | 1557.93 ± 89.54 | 49 | - | 0.424 |
| Time with aggressive colour on (s)* | 8 | 1775.38 ± 32.48 | 10 | 1708.98 ± 124.58 | 49 | - | 0.424 |
| Frequency of caudal swings* | 8 | 4.625 ± 5.527 | 10 | 26 ± 33.269 | 25 | - | 0.174 |
| Frequency of charges | 8 | 4.875 ± 9.342 | 10 | 22.5 ± 28.96 | 1.645 | 16 | 0.119 |
| Frequency of bites | 8 | 65.875 ± 67.397 | 10 | 67.3 ± 110.924 | 0.032 | 16 | 0.975 |

\*, differences tested with the Mann-Whitney U test.

Interactive live and mirror fights in this species follow a highly stereotyped sequence, starting with threat displays (opening of opercula, distension of fins) and switching to overt aggression (charges, bites) after a few minutes [11,14]. We investigated whether this also occurred in non-interactive fights and if it could be influenced by the type of stimuli. A two-way repeated measures ANOVA on the time with open opercula, a threat behaviour, showed that it was higher in the first half, regardless of stimulus type (within-effects combat phase, F_(1,16)_ = 19.782, p < 0.001; between-effects stimulus type, F_(1,16)_ = 0.036, p = 0.852; effects interaction, F_(1,16)_ = 0.150, p = 0.704; Fig. 2). Attack behaviours were more frequent in the second half of the fight, and again no effect of stimulus type was recorded (bites, within-effects combat phase, F_(1,16)_ = 5.960, p = 0.027; between-effects stimulus type, F_(1,16)_ = 0.001, p = 0.975; effects interaction, F_(1,16)_ = 0.590, p = 0.454; Fig. 2). Air breathing, an indicator of metabolic activity [27], was higher in the second half of the test, independently of stimulus type (within-effects combat phase, F_(1,16)_ = 9.991, p = 0.006; between-effects stimulus type, F_(1,16)_ = 0.726, p = 0.407; effects interaction, F_(1,16)_ = 2.259, p = 0.152; Fig. 2). Accordingly, the frequency of air breathing correlated with the frequency of attack behaviours (bites, r = 0.537, N = 18, p = 0.019) but not with the duration of threat displays (opening of opercula, r = 0.240, N = 18, p = 0.338).

**Figure 2.**
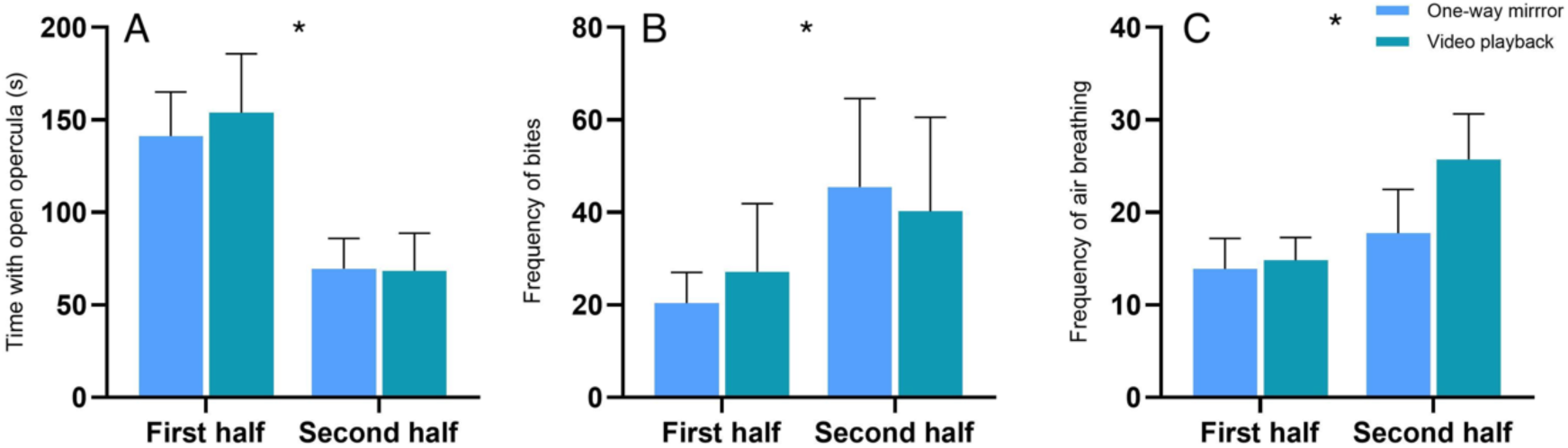
(A) Duration of open opercula, (B) frequency of bites and (C) frequency of air breathing in the first (0-15 min) and second (15-30 min) half of the aggression trials. Focal fish were presented with a conspecific fighting a one-way mirror, live (One-way mirror) or as video images (Video playback). Mean ± S.E. are shown. * represent significant differences (p < 0.027) between the first and second half of the trial. The live and video stimuli triggered similar responses in the focal fish (p > 0.407).

### Hormones

There was a marked increase in plasma KT levels after the 30 min aggression challenge. The one-way mirror and video playback aggression stimuli resulted in an average KT increase of 4.2 and 5-fold, respectively, as compared to control (one-way ANOVA on KT log-transformed values, F(2,28) = 10.189, p < 0.001; Fig. 3). Post-hoc comparisons confirmed a significant difference of both the one-way mirror (p = 0.003) and video playback (p = 0.002) groups to the control group, while there was no difference between fish from the two aggression treatments (p = 0.999). Post-fight KT levels of fish from the aggression-elicited groups did not correlate with aggression or activity variables displayed during fights (p > 0.095).

**Figure 3.**
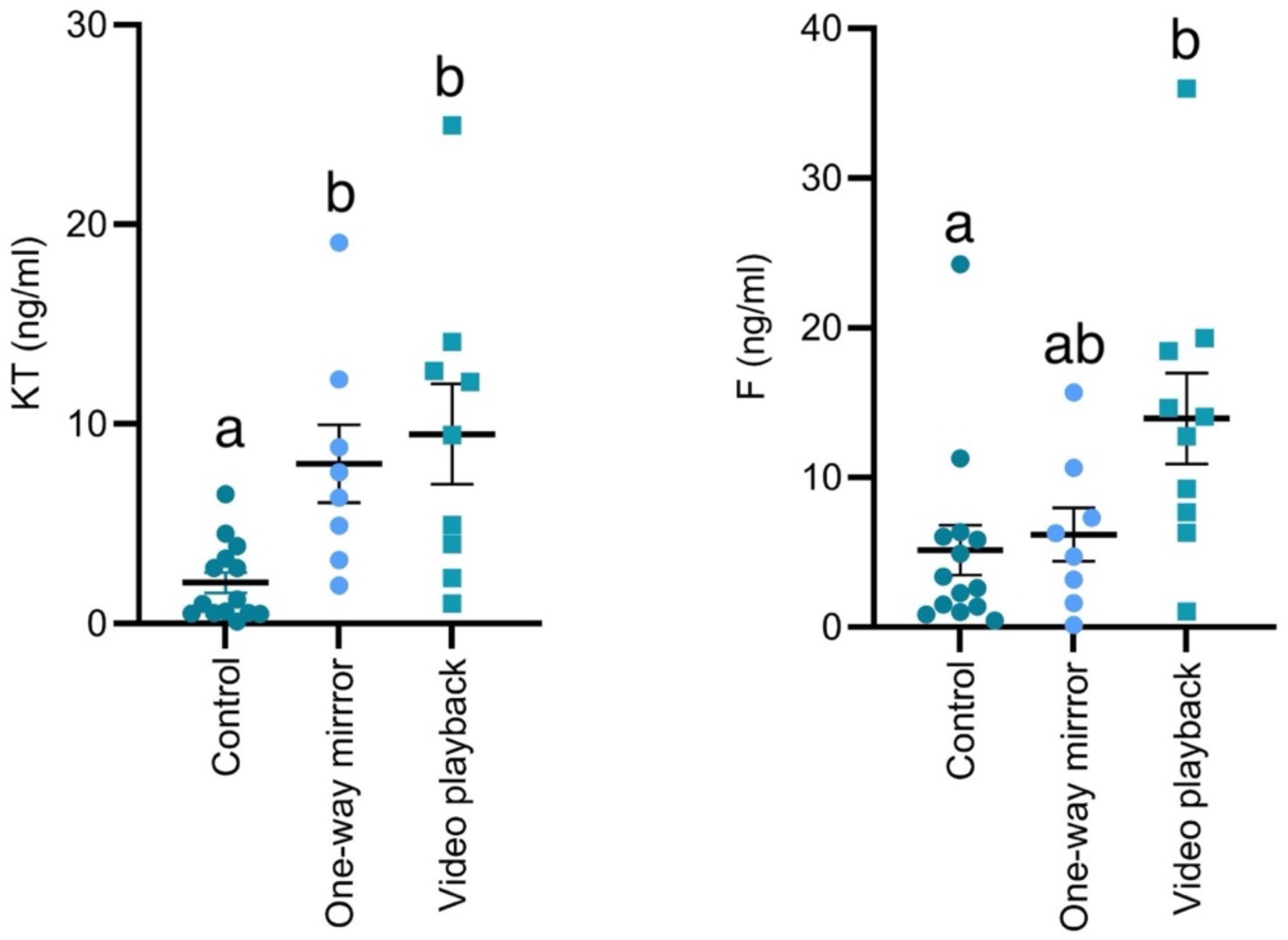
Post-test plasma levels of 11-ketotestosterone (KT) and cortisol (F) in fish presented for 30 min with a conspecific fighting a one-way mirror, live (One-way mirror) or as video images (Video playback). Control fish were presented with either an empty tank or with a video playback of the empty tank. Mean ± S.E. are shown. Different letters represent significant differences (p < 0.05) between groups.

Levels of F also differed between treatments (F(2,29) = 4.721, p = 0.017; Fig. 3). When presented with video images of a conspecific, F levels increased almost 3-fold as compared to controls (p = 0.017). This increase was less pronounced in fish presented with the live conspecific, with post-hoc comparisons with the control (p = 0.943) and video playback group (p = 0.077) not being significantly different. Again, no significant correlation with fight and activity variables could be detected (p > 0.094). Levels of F and KT were also uncorrelated (r = 0.136, N = 31, p = 0.465).

### Video playback vs one-way mirror

To further test if the behavioural and physiological response to the live and video stimuli were comparable, a PCA including all behavioural variables and post-fight KT and F levels was performed. The first three factors of the PCA explained 74.8 % of the variance (Supplementary Table 1). Fish from the control group formed a separate cluster but there was significant overlap between the fish exposed to the live or video playback stimuli (Fig. 4). Still, variability of the response was apparently higher for the video playback group as it showed a larger dispersion of individuals to the group centroid (average distance to centroid: control = 0.5745; one-way mirror = 1.0866; video playback = 2.6366).

**Fig. 4.**
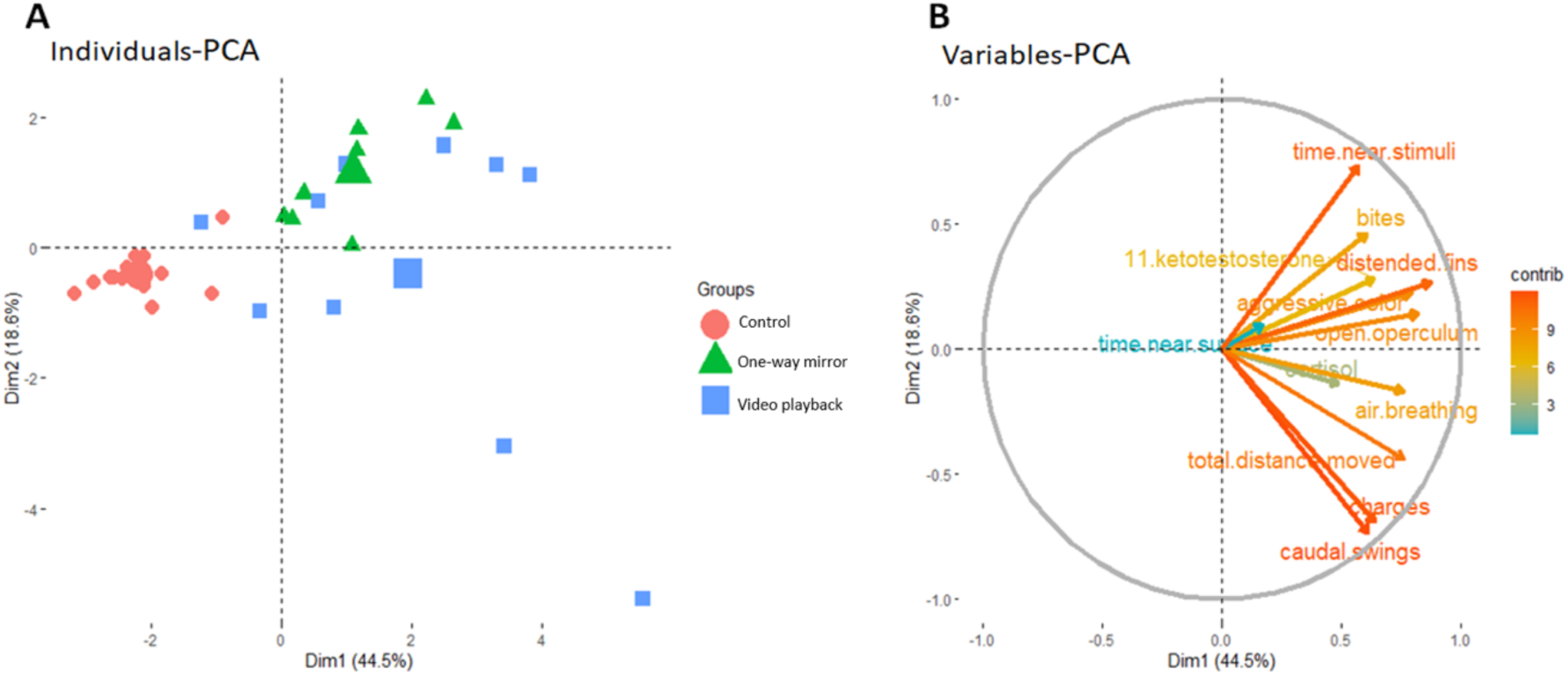
Two first components of a Principal Component Analysis with all the measured endocrine and behavioural variables. (A) individual data; large symbols represent the centroid of the groups; (B) variable loadings; the direction of arrows represents the correlation of variables and size their relative contribution. Numbers in the axis’s legend represent the percentage of variation explained by each component.

## Discussion

### Video playback as a tool to study fish aggression

Results show for the first time in a fish a robust endocrine response to aggression elicited by video images of a conspecific. Plasma levels of KT, the most potent androgen in fish [28], increased over 5-fold after observing the opponent on screen and this increase was comparable to the response observed towards a live conspecific fighting behind a one-way mirror. Cortisol also increased significantly after the video playback fight, as previously reported for mirror or paired fights in *B. splendens* [11,29], while differences to the live stimuli were not significant.

This is also the first study in fish comparing responses to a conspecific fighting behind a one-way mirror, presented live or through video playback. Previous studies in fish have validated video playback as a technique by comparing a non-interactive video stimulus with a live interactive stimulus, not allowing to disentangle the two factors (e.g., [30]). This may have contributed to a mismatch in the response of focal animals towards live and video stimuli in some studies (e.g., [31]). The behavioural response of *B. splendens* males to the video playback was evident and comparable to the response towards the live opponent. The fight followed a similar sequence towards live and video stimuli, starting with threat displays and progressing to overt attacks, as previously described [11,14]. A PCA combining all the behavioural and endocrine data corroborated these findings as there was extensive overlap in the distribution of fish from the one-way mirror and video playback groups. Nevertheless, data dispersion seemed to be higher for fish in the video playback group. The reasons for this are not clear but may relate with variation between animals in the sensorial perception of video images (for a discussion see [32]). A few studies have used video playback as a tool in this species. Allen and Nicolleto [20] showed that males responded aggressively to manipulated video images of conspecific males of different fin sizes by opening the opercula, the first element to be displayed in a fight [31], and by displaying other aggression elements. Similar results were obtained by Neri [34], with males orienting towards a video playback of a conspecific and opening the opercula, although some fish showed only a weak response, corroborating our observation that video images produced more variable results than the live stimulus. Clotfelter and collaborators [35] show that males and females chose between pairs of computer-generated stimuli differing in size or display features, as assessed by time spent close to each stimulus, but the frequency or duration of aggressive displays was not reported. None of these previous studies, however, compared the response of *B. splendens* to video playback with an equivalent live stimulus nor provided physiological measures of this response.

If properly validated, video playback provides a powerful tool for the study of behaviour. This technique allows the presentation of the same invariant stimulus to animals from different treatments or along different periods of time, minimizing variation across trials. Further, digitization and animation techniques allow precise control that can be used to produce synthetic animations to test the response to particular features (e.g., [30,31]), or to design interactive video playback experiments where the stimulus output depends on the behaviour of the focal animal (e.g., [38]). One relevant question is whether measuring the response towards a non-interactive stimulus, either video or live, is appropriate [36]. Real-life fights are, by nature, interactive and the absence of dynamic feedback from the opponent, and the impossibility to display the full suite of aggressive behaviours (e.g., circling or chases during fights), may generate a poor response. Although this study was not designed to specifically test the importance of fight interaction for aggressive displays, results can be compared with previously published data from our lab where the aggressive response of males of the same strain, and under similar experimental conditions, were tested with matched for size opponents behind a transparent partition [11]. While controls in both studies had a similar frequency of air breathing, interactive fights caused a 2.5 to 3 - fold higher frequency of this behaviour as compared with one-way mirror fights, suggesting that they are metabolically more demanding (compare Table 1 in this study with Table 2 in [11]). The duration of threat displays (average time with opercula open, 211 s vs 372 s) and of attacks (average frequency of bites, 66 vs 95) was also higher when dynamic feedback was available (compare Table 2 in this study with Table 3 in [11]). Accordingly, average post-fight plasma levels of KT (8.0 vs 10.6 ng/ml) and F (6.2 vs 13.0 ng/ml) were higher when both males could see each other (compare Figure 3 in this study with Figure 5 in [11]). Although this comparison supports the hypothesis that aggressive behaviour, and associated physiological responses, is stronger in interactive fights, results should be interpreted with caution as these were separate experiments and uncontrolled differences between studies may explain the results. Even if the behavioural and endocrine output is reduced towards non-interactive stimuli, we argue that video playback, which allows a full standardization of stimuli, is a useful tool to measure variation in aggression levels across groups in *B. splendens*.

### Hormones and fish aggression

Post-fight plasma androgen levels were markedly elevated in fish responding to the aggression challenge, regardless of stimulus type. These results corroborate previous data from our lab, both for KT and for testosterone (T), using as stimuli mirrors and a conspecific behind a transparent partition [11]. Similar findings have been reported for other strains of *B. splendens*, including wild-type [29], as well as for the cichlid *Astatotilapia burtoni* [23] and *Pundamilia* spec. [37]. These results are consistent with the challenge hypothesis, which stipulates that males respond to aggression challenges with a rapid increase in androgen levels [4]. Indeed, it has been shown that observing other males fighting is enough to increase androgen levels in a bystander [39]. Although a lack of response (e.g., [40–42]) or even a decrease (e.g.,[43]) in androgen levels after unsolved fights has been reported, a recent meta-analysis confirmed that androgen levels increase in response to social challenges in teleost fish [44].

If it is becoming increasingly evident that androgen levels increase above reproductive levels when animals are socially challenged, the functional role of this increase is still debatable. If its function is to facilitate the expression of aggressive behaviour to better defend reproductive resources [4], a co-variation in androgen levels and aggression is expected. However, there was no correlation between post-fight androgen levels and aggressive behaviour in this study, in agreement with several other works in fish (e.g., [11,45]). Further, while T administration has been shown to induce a moderate increase in the duration of opercula opening in *B. splendens* [46], castration, expected to suppress androgen secretion, failed to inhibit aggressive behaviour in this species [7] and also in the Mozambique tilapia *Oreochromis mossambicus* [6]. One hypothesis that remains to be tested, is whether the androgen increase may relate to sustaining fight behaviour. Although there is no information available for the duration of *B. splendens* fights in nature, paired fights staged by breeders in Asia can take several hours to be settled (personal observations conducted in fish fighting rings in Thailand). Androgens have been shown to increase in humans after physical activity and induce insulin-related effects in skeletal muscles by the upregulation of glucose transporter type 4 [47]. It is thus possible that the observed increase in KT is associated with preparing the muscles for sustained activity or recovery [48]. This hypothesis can be tested by exposing the fish to prolonged fights (ideally using standardized stimuli, as video playback), while manipulating the androgen system. Alternatively, the androgen increase may prepare fish not for the present but for future fights. Indeed, androgens have been proposed to mediate the winner effect, whereby winning a fight increases the probability of winning a future challenge [49]. This seems to be corroborated by a study in *O. mossambicus* where blocking androgen receptors cancelled the winner effect [50] and by a study in killifish *Kryptolebias marmoratus* where pre-fight KT levels correlated with aggression and winning probability [51]. Interestingly, winning is not always needed to trigger the winner effect as fish fighting their mirror image, where there is no winner experience, may also increase the probability of winning future fights [37]. Thus, it seems possible that the androgen increase after a fight experience may underlie the observed winner effect. However, evidence has been accumulating that the loser effect may be mediated by a different physiological mechanism. Winners and losers have been shown to have a similar increase in post-fight androgen levels (e.g., [52]; unpublished data for *B. splendens*) and androgen administration to losers failed to reverse the loser effect [50]. Losing also seems to be a more salient experience in zebrafish as losers show a stronger change in brain transcriptome as compared with winners, and different genomic pathways were implicated in the winner and loser experience [53]. It is thus possible that losing a fight overturns the positive impact of androgens in aggression, being mediated by other mechanisms, either central or peripheral. Manipulating video playback during an ongoing fight to potentially modulate the perception of winning/losing would be a useful tool to confirm this hypothesis in fish.

Plasma F levels also increased in focal fish fighting the video playback aggression stimulus, in agreement with previous studies in this [11] and other (e.g., [12]) fish species, but not when fighting the conspecific behind the one-way mirror. Interestingly, the response to a mirror challenge of males from the two parental strains (one fighter and one wild-type) that originated the animals used in this study was consistent for KT but not for F, with F increasing in wild-type but not in fighter males [29]. It thus seems possible that the difference in the F response to the two types of stimuli could result from spurious genetic variation between fish tested with the video playback and with the live stimulus. Variability in the F response after exposure to a stressor between a wild-type and a fighter strain, different than those used at our lab, had also been previously reported [54].

These results would seem to suggest that F correlates positively with aggressive behaviour in fish. However, a number of studies have shown opposite results. For example, in lines of rainbow trout *Oncorhynchus mykiss* selected for low and high cortisol responsiveness to a stressor, low responders were more frequently associated with a dominant status [55] and were more aggressive [56] than high responders. One possibility is that short-term F responsiveness is unrelated to aggression but long-term F profile is. First, post-fight F levels did not correlate with aggressive behaviour in this study (see also [11]) but wild-type fish, which are less aggressive than domesticated fighters [15,57], have consistently elevated F levels [29]. Second, in subordinate and dominant rainbow trout, F increases rapidly in both participants in a fight, but it starts decreasing in the dominant fish while it continues to increase in the subordinate [58]. Third, in the same species, short-term F exposure did not influence aggression while long-term treatment inhibited it [59]. Thus, while short-term increase in F during fights may relate to its role in glucose-regulation and glycogen-repletion processes [60], dominance and enhanced aggression may be associated with chronically low F levels. This hypothesis may be tested by measuring the impact in aggressive behaviour of short and long-term manipulation of the HPI axis. Again, video playback can be a useful tool to investigate this topic in *B. splendens*.

### Conclusions

In conclusion, video playback offers an excellent tool to study under laboratory settings the intrinsic motivation for aggression in *B. splendens*. The species is a promising model for decoding the role of steroid hormones in aggressive behaviour and recent developments, in particular the sequencing of its genome [61], will allow probing in more detail the physiological pathways underlying variation in aggression in this interesting fish.

## Supplementary Material

**Supplementary table 1.** Loadings for the three first factors of a Principal Component Analysis (PCA) and percentage of variance explained by each factor. The PCA included all behavioural and endocrine data of fish presented with an empty tank or white screen (control), opponent behind a one-way mirror in an adjacent tank, and video playback of an opponent fighting a one-way mirror.

|  | PC1 | PC2 | PC3 |
| --- | --- | --- | --- |
| 11.ketotestosterone | 0.275 | 0.189 | -0.146 |
| Cortisol | 0.21 | -0.093 | 0.246 |
| Aggressive colour | 0.343 | 0.15 | -0.283 |
| Air breathing | 0.328 | -0.113 | 0.393 |
| Bites | 0.261 | 0.309 | 0.338 |
| Caudal swings | 0.263 | -0.491 | -0.1 |
| Charges | 0.275 | -0.462 | -0.123 |
| Distended fins | 0.377 | 0.178 | -0.226 |
| Open operculum | 0.355 | 0.094 | -0.176 |
| Total distance moved | 0.329 | -0.294 | 0.121 |
| Time near surface | 0.073 | 0.063 | 0.668 |
| Time near stimuli | 0.247 | 0.489 | -0.047 |
| Percentage of variance explained | 44.5 | 18.6 | 11.7 |

Supplementary video 1. Example of the response of a focal fish to the video playback. 30 s pre-stimulus (white screen) and the first 60s of the fight against the video of a conspecific fighting a one-way mirror is shown.

